# Cholinergic waves have a modest influence on the transcriptome of retinal ganglion cells

**DOI:** 10.1101/2024.12.05.627027

**Authors:** Rachana Deven Somaiya, Matthew A. Po, Marla B. Feller, Karthik Shekhar

## Abstract

In the early stages of development, correlated activity known as retinal waves causes periodic depolarizations of retinal ganglion cells (RGCs). The β2KO mouse, which lacks the β2 subunit of the nicotinic acetylcholine receptor, serves as a model for understanding the role of these cholinergic waves. β2KO mice have disruptions in several developmental processes of the visual system, including reduced retinotopic and eye-specific refinement of RGC axonal projections to their primary brain targets and an impact on the retinal circuits underlying direction selectivity. However, the effects of this mutation on gene expression in individual functional RGC types remain unclear. Here, we performed single-cell RNA sequencing on RGCs isolated at the end of the first postnatal week from wild-type and β2KO mice. We found that in β2KO mice, the molecular programs governing RGC differentiation were not impacted and the magnitude of transcriptional changes was modest compared to those observed during two days of normal postnatal maturation. This contrasts with the substantial transcriptomic changes seen in downstream visual system areas under wave disruption in recent studies. However, we identified ∼238 genes whose expression was altered in a type-specific manner. We confirmed this result via *in situ* hybridization and whole-cell recording by focusing on one of the downregulated genes in aRGCs, *Kcnk9*, which encodes the two-pore domain leak potassium channel TASK3. Our study reveals a limited transcriptomic impact of cholinergic signaling in the retina and instead of affecting all RGCs uniformly, these waves show subtle modulation of molecular programs in a type-specific manner.

**SIGNIFICANCE STATEMENT:** Spontaneous retinal waves are critical for the development of the mammalian visual system. However, their role in transcriptional regulation in the retina across the diverse retinal ganglion cell (RGC) types that underpin the detection and transmission of visual features is unclear. Using single-cell RNA sequencing, we analyzed RGC transcriptome from wild-type mice and mice with disrupted retinal waves. We identified several genes that show RGC-type-specific regulation in their expression, including multiple neuropeptides and ion channels. However, wave-dependent changes in the transcriptome were more subtle than developmental changes, indicating that spontaneous activity-dependent molecular changes in retinal ganglion cells are not primarily manifested at the transcriptomic level.

## Introduction

The visual system is a classical model for exploring the role of neuronal signaling in the development and refinement of neural circuits (Stevens-Sostre and Hoon, 2024). Retinal ganglion cells (RGCs), which are the sole output neurons of the retina, comprise more than 40 distinct types in mice that can be classified using morphological, physiological, connectomic, and molecular attributes (Baden et al., 2016; Bae et al., 2018; Rheaume et al., 2018; Tran et al., 2019; Goetz et al., 2022; Huang et al., 2022). In adult mice, RGCs of a given type have highly precise connections with other types of presynaptic neurons in the inner retina, which renders most individual RGC types selectively responsive to a small subset of visual features such as orientation, directional motion, and edges (Sanes and Masland, 2015; Kerschensteiner, 2022).

During development, as RGCs differentiate into discrete types, they exhibit spontaneous wave-like activity, known as retinal waves, which precedes photoreceptor-driven responses (Galli and Maffei, 1988; Meister et al., 1991; Wong, 1999). Retinal waves span a protracted period during early development, starting on E17 and persisting through eye-opening on postnatal day 12 (P12) (Blankenship and Feller, 2010). These waves travel throughout the visual system and are the primary source of activity in the dorsal lateral geniculate nucleus (dLGN) (Grubb et al., 2003; Murata and Colonnese, 2016), superior colliculus (SC), and primary visual cortex (V1) prior to the onset of vision (Ackman et al., 2012; Burbridge et al., 2024).

In the retinofugal targets, numerous studies highlight the critical role for cholinergic waves in shaping the development of visual circuits. Genetic or pharmacological blockade of these waves has been shown to reduce the segregation of eye-specific RGC terminals in the dLGN and SC and retinotopic refinement of RGC axons in the SC (reviewed in Huberman et al., 2008; Blankenship and Feller, 2010; Kirkby et al., 2013; Kerschensteiner, 2014; Arroyo and Feller, 2016; Choi et al., 2021). More recently, several studies have revealed that perturbations in perinatal cholinergic wave activity leads to significant transcriptomic changes in downstream visual centers such as in the neurons of the SC and V1 (Guillamón-Vivancos et al., 2022; Burbridge et al., 2024).

How disruption of spontaneous activity impacts the transcriptome at the level of RGCs is not as well studied. A recent study showed that reducing the frequency of cholinergic waves induces widespread transcriptomic changes in RGCs, as assayed by bulk RNA sequencing (bulk RNAseq) (Negueruela et al., 2024). However the impact on individual RGC types is unknown. This is critical because many studies have shown that waves influence the development of dendrites in specific subsets of RGCs (Sernagor and Grzywacz, 1996; Bansal et al., 2000; Sernagor and Mehta, 2001; Mehta and Sernagor, 2006; Liu et al., 2009; Xu et al., 2010; Elias et al., 2018; reviewed in Tian, 2008) as well as the establishment of the functional circuitry corresponding to a subset of RGCs that selectively respond to horizontal directional motion in the mouse retina (Tiriac et al., 2022). Together, these studies suggest that early spontaneous activity likely regulates factors through RGC type-dependent manner to influence the synapse-specific wiring.

To reveal whether the impact of spontaneous activity on different RGC types is manifested at the level of gene expression patterns, we used single-cell RNA sequencing (scRNAseq) to compare RGC transcriptome between wild-type (WT) and mice lacking the β2 subunit of neuronal nicotinic receptor. Genetic removal of the β2 subunit of the nicotinic acetylcholine receptor (nAChR) inhibits cholinergic signaling, thereby disrupting retinal waves between E16 and P7 (Bansal et al., 2000; Rossi et al., 2001; Burbridge et al., 2014; Voufo et al., 2023). This mouse model, henceforth called β2KO, has been used widely to study the developmental roles of spontaneous activity-induced by cholinergic waves. Our analysis focused on several key aspects: the effects of these waves on RGC type diversification, the global transcriptomic shifts across all RGC types, and the type-specific changes to reveal whether certain RGC types have higher sensitivity to wave disruption.

## Methods

### Mice

All procedures were approved by University of California, Berkeley Institutional Animal Care and Use committee and were in accordance with the National Institutes of Health Guide for the Care and Use of Laboratory Animals, the Public Health Service Policy, and the Society for Neuroscience Policy on the Use of Animals in Neuroscience Research. Mice used in this study were P7-P60 and of both sexes. Our control experiments were performed using C57BL/6J (RRID:IMSR_JAX:000664). β2KO mice that were deficient in the beta-2 subunit of the nicotinic acetylcholine receptor were originally provided by Dr. Art Beaudet from Baylor College of Medicine (Xu et al., 1999) and maintained in our laboratory onto a C57BL/6J background. Genotyping was performed by Transnetyx.

### RGC isolation for scRNAseq and bulk RNAseq

P7 C57BL/6J and β2KO mice were anesthetized in an isoflurane induction chamber and euthanized by decapitation. For both scRNAseq and bulk RNAseq, at least 3 biological replicates (each referred to as a sample) were run for each genotype and each sample included RGCs pooled from 3-4 mice. All the solutions were filtered using 0.22 µm syringe filter (Fisher #9720511). Eyes were enucleated and retinas were immediately dissected in oxygenated (95% oxygen/5% carbon dioxide) Ames’ media with L-glutamine and sodium bicarbonate (pH ∼7.4) at room temperature under light scope. Retinas (6 to 8 per sample) were digested for 10 min at 37°C in 70 µl papain (Fisher #LS003126) + 50 µl DNase (Sigma #D4527-40KU) + 50 µl L-cysteine (Sigma #C1276-10G) in 5 ml Ames, which was activated at 37°C for 15 min. Samples were gently swirled once during this incubation. Following this, papain was removed and samples was gently triturated with P1000 pipette tips in 1 ml 10x ovomucoid (Fisher #NC9931428) + 50 µl DNase + 9 ml Ames. Manual trituration was performed using 2 ml of ovo solution at a time and the supernatant was filtered using 40 μm cell strainer in a new tube. This cell suspension was spun down at 450 *g* for 8 min. Supernatant was discarded and the pellet was resuspended in 250 μl of Ames + bovine serum albumin (BSA). This solution was prepared using 27 ml Ames + 3 ml 4% BSA (Sigma #A9418-5G) + 50 µl DNase. Cells were counted using hemocytometer (1:100 dilution factor with trypan blue). 10 million cells were incubated in 10 µl CD90.2 microbeads (Miltenyi Biotec #130-121-278) for 10 min at 4°C. Following incubation, 6 ml Ames + BSA solution was added and cells were spun down at 450 *g* for 8 min. Supernatant was discarded and cells were resuspended in fresh 500 µl Ames + BSA solution. Cell suspension was subjected to magnetic-activated cell sorting (MACS) using large cell columns (Miltenyi Biotec #130-042-202) to sort RGCs. To obtain a high number of RGCs, the tube containing the cells was washed with 500 µl Ames + BSA solution for 2 times. To obtain better purity, MACS was repeated one more time on eluted CD90.2^+^ RGCs using fresh columns. This cell suspension was spun down at 450 *g* for 8 min and the supernatant was removed. Pellet obtained was resuspended in ∼1000 μl of Ames + BSA and cell concentration was counted using hemocytometer (dilution factor 2). This spun down strategy was employed to obtain a highly concentrated sample in a small volume of ∼100 μl Ames + BSA collected in a Lo-bind tube.

### RNA extraction for bulk RNAseq

All steps were performed using manufacturer’s protocol for Direct-zol™ RNA Miniprep Plus with TRI Reagent® (VWR International #76020-112). Briefly, ∼100 μl Ames + BSA of RGC suspension was mixed with 300 μl of Trizol by pipetting up and down. 300 μl of 100% ethanol was added and mixed by pipetting. This mixture was transferred into the Zymo-Spin column in a collection tube and centrifugation was performed (all centrifuge steps were performed at 15000 rcf for 30 sec unless mentioned otherwise). The flow-through was discarded. The column was then transferred into a new collection tube and 400 μl RNA wash buffer was added. While the sample was spinning down, in a RNase-free tube, 5 μl of DNase was combined with 75 μl of DNA digestion buffer and mixed by gentle inversion. This mixture was added to the column and incubated for 15 min at room temperature. Following incubation, 400 μl RNA prewash was added to the column, centrifugation was performed, and flow-through was discarded. This step was repeated once. To the column, 700 μl RNA wash buffer was added and the tube was spun down for 1 min. The column was then transferred carefully to a DNA Lo-bind tube, 50 μl of DNase/RNase-free water was added, and followed by centrifugation. Sample was frozen in −80°C until bulk RNAseq was performed.

### Library preparation and sequencing for bulk RNAseq

The quality of total RNA and poly-dT-enriched mRNA was evaluated using an Agilent 2100 Bioanalyzer. Libraries were prepared with the KAPA mRNA HyperPrep Kit (Roche KK8581). Truncated universal stub adapters were ligated to cDNA fragments, which were then amplified into full-length Illumina adapters through 14 cycles of PCR using unique dual-indexing primers. Library quality was assessed on an AATI (now Agilent) Fragment Analyzer, and library molarity was quantified using qPCR with the KAPA Library Quantification Kit (Roche KK4824) on a Bio-Rad CFX Connect thermal cycler. These libraries were then pooled by molarity, followed by sequencing on an Illumina NovaSeq X system for 2 × 150 cycles to target at least 25 million reads per sample. Sequencing for all the samples was performed together to avoid any run-to-run bias. Using Illumina BCL Convert v4 with default settings on a server running CentOS Linux 7, FASTQ files were generated and demultiplexed.

### 10X genomics for scRNAseq

Single-cell libraries on RGCs isolated were prepared using Chromium Next GEM Single Cell 3’ Kit v3.1 (10X Genomics, Pleasanton, CA) following manufacturer’s protocol. On average, for each sample, cells were loaded to ultimately recover 5000-10000 cells. At least 3 biological replicates (each referred to as a sample) were run for each genotype and each sample included RGCs pooled from 3-4 mice. All 10X-related steps were performed by QB3 Genomics of University of California, Berkeley, Berkeley, California, RRID: SCR_022170. Libraries were sequenced on the Illumina NovaSeq platform at the University of California, Irvine Genomics Research & Technology Hub.

### In situ RNA hybridization (ISH) and immunohistochemistry (IHC)

Retinas were dissected as described above, fixed in 4% paraformaldehyde overnight at 4°C, and then submerged in 30% sucrose solution for ∼3 days at 4°C. Using a cryostat, retinas were thinly sliced to a thickness of 16 µm.

RNAscope® Multiplex Fluorescent Detection Kit v2 (ACD #323110) was performed to detect *Kcnk9* mRNA (ACD #475681) using the manufacturer’s instructions for fixed frozen tissue. Only few changes were made to their established protocol. These were: incubating tissue in 4% PFA for 1 hr instead of 15 min; 5 min of target retrieval; and protease treatment for only 15 min. Opal dyes (Akoya #FP1487001KT) at 1:750 dilution were used for signal detection. Following ISH, IHC was performed to detect SMI32^+^ (1:250 dilution, BioLegend #801702,) and RBPMS^+^ (1:500 dilution, Abcam #ab152101) cells. Blocking solution used was 1% BSA + 0.2% Triton X-100 in 1X Tris-buffered saline (TBS). Retinal slices were blocked for 30 mins at room temperature and incubated in primary antibodies (prepared in blocking solution) overnight at 4°C. Next day, retinal slices were washed for 3 times (15 min each) in 0.1% Tween 20 and 0.1% Triton X-100 in 1X TBS solution and incubated in secondary antibodies (1:500 dilution prepared in blocking solution, Jackson ImmunoResearch #111-605-003, Fisher #A28175) for 2 hr at room temperature. After a few washes, slides were mounted with media containing DAPI (Fisher #H-1500).

Retinal slices were imaged on a Zeiss LSM880 Laser Scanning Confocal Microscope with a Plan-Apochromat 20x/0.8 M27 objective and resolution of 2048 by 2048 pixel (pixel size: 0.208 µm^2^) at Molecular Imaging Center at University of California, Berkeley. Z-stacks with 2 µm were used to acquire images. Representative images in the figures are maximum intensity projections, while colocalization of *Kcnk9* mRNA signal with RBPMS and SMI32 was confirmed using single-plane images.

### Quantification and statistical analysis of ISH and IHC signal

RGCs were identified in each image to allow for reporting of per-cell metrics. Cells were segmented on the RBPMS IHC channel using the pretrained Cellpose model cyto3, which was fine-tuned using manually annotated images (Stringer et al., 2021; Pachitariu and Stringer, 2022). DAPI stains were also provided to the segmentation model to distinguish proximal cells. RGCs identified through this segmentation that colocalized with SMI32 were labeled as αRGCs.

Quantification of the ISH signal was performed using the Python package Big-FISH (Imbert et al., 2022). Signal was quantified using the number of puncta per cell and average intensity per cell. To count the number of *Kcnk9* puncta in each cell, the image was first background subtracted using a mean subtraction within a 2-pixel-wide square kernel. A Laplacian of Gaussian filter was then used to denoise the image. Puncta were identified as local maxima in the filtered image. Finally, an intensity threshold determined through Otsu’s method was used to filter for high-intensity puncta (Otsu, 1979). A mask from the cell segmentation was used to then compute the number of puncta per cell. Average intensity per cell was computed by averaging the intensities of all pixels within a cell in the Cellpose mask.

Significance testing between conditions for both puncta counts and intensities was performed using a Mann-Whitney U-Test, taking each cell as an individual observation.

### Patch-clamp electrophysiology of RGCs

Voltage-clamp experiments on αRGCs to identify leak channel-associated conductance were performed as described previously (Wen et al., 2022). P7-P12 (all before eye opening) C57BL/6J and β2KO mice were anesthetized in an isoflurane induction chamber and euthanized by decapitation. Eyes were enucleated and retinas were immediately dissected at room temperature under light scope in oxygenated (95% oxygen/5% carbon dioxide) Ames’ media with L-glutamine and sodium bicarbonate with pH of 7.4, adjusted using NaOH/HCl (∼300 mOsm). Isolated retinas were placed flat in the recording chamber with a continuous perfusion of oxygenated Ames at ∼2 ml/min. A temperature of 32–34°C was maintained for the duration of experiments. Recordings were performed in light, except when taking morphology images under two-photon illumination.

MultiClamp 700B amplifier, Digidata 1440A digitizer, and pCLAMP 10.7 software were used for patch-clamp experiments. Patch pipettes were pulled from borosilicate glass capillaries (with an outer diameter of 1.5 mm and an inner diameter of 1.1 mm) and resistance of ∼5 MΩ (PC-10 pipette puller; Narishige). The internal solution contained (in mM): 125 K-gluconate, 2 CaCl_2_, 2 MgCl_2_, 10 EGTA, 10 Hepes, 0.5 Mg–ATP, and 2 Na_2_–GTP (∼290 mOsm). To fill the cell, 10 µM Alexa Fluor 488 hydrazide (Life Technologies #A10436) was also added to the internal solution. pH of the solution was ∼7.2 and adjusted using KOH. The liquid junction potential corrected was −13 mV. αRGCs were identified based on their large soma size and dendritic field, which was identified via the Alexa Fluor 488 dye-filled cell (Krieger et al., 2017). Cells were voltage-clamped at −73 mV for 100 ms, following holding at +17 mV for 500 ms and ramped to −133 mV within 500 ms. In bath solution, tetrodotoxin (TTX, to block voltage-gated sodium channels) at 500 nM and 4-Aminopyridine (4-AP, to block voltage-gated potassium channel) at 5 mM were added to Ames to isolate leak channel-associated conductance.

Current required to hold WT and β2KO RGCs at +17 mV was calculated by subtracting current required to hold at −73 mV from current required to hold at +17 mV during ramp protocol. Statistical analysis was performed using R. Data in the figure 4G is represented as means with standard deviation. Welch two-sample t-test was performed as unequal sample size and without assuming equal variances between the samples. P value less than 0.05 was defined as significant.

### Code availability

Jupyter notebooks required to reproduce the analyses presented here are available on our GitHub page: https://github.com/shekharlab/WaveRGCs.

### Data availability

All raw and processed scRNAseq datasets reported in this study will be made publicly available via NCBI’s Gene Expression Omnibus (GEO) Accession Number GSE283463. Processed h5ad files are available at https://github.com/shekharlab/WaveRGCs. These h5ad files contain all relevant metadata and log-normalized counts.

### Alignment and quantification of scRNAseq data

Sequencing read libraries generated in this work (P7 WT and β2KO) were aligned to the UCSC mm10 genome assembly using Cellranger v7.1.0 (10X Genomics). This alignment was performed separately on each sample (three P7 WT samples, four P7 β2KO samples), and the resulting cells-by-genes expression matrices were combined to obtain an expression matrix for each condition.

Preprocessing of expression data was performed using the Scanpy workflow (Wolf et al., 2018). Cells expressing less than 700 genes, containing less than 5,000 transcripts, or with over 10% of its counts from mitochondrial genes were filtered out of the data. The count matrix **X** was then normalized such that each column sums to 10,000 (corresponding to total reads per cell) and logarithmized such that, for a normalized matrix entry *x_ij_*, its corresponding entry in the log-transformed matrix is *log* (*x_ij_* +1).

### Clustering

Highly variable genes (HVGs) were identified using a previously published method (Satija et al., 2015) based on the ratio of a gene’s expression variance to its mean. Briefly, the column mean (µ*_j_*) and variance (σ*_j_* ^2^) of **X** were computed and used to calculate a dispersion metric

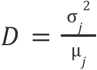

which was used to split all genes into twenty bins. Within each bin, dispersion metrics were z-scored, and those genes with a z-score greater than 2 were identified as highly variable. The expression matrix was then subset to HVGs and scaled along columns (genes) to unit variance and zero mean. Principal components analysis (PCA) was applied to reduce the dimensionality of the data, then batch correction was applied via Harmony (Korsunsky et al., 2019). The top 30 batch-corrected PCs were used to build a k-nearest neighbors graph, which was used to cluster cells via the Leiden algorithm (Traag et al., 2019) and create a Uniform Manifold Approximation and Projection (UMAP) embedding of cells for visualization (McInnes et al., 2020). We emphasize that UMAP was used only for the post hoc visualization of clusters, and was not used to guide the graph construction or clustering procedure.

### Type identification

P7 WT and β2KO clusters were assigned to classes based on expression of known retinal cell class markers. Clusters expressing *Slc17a6*, *Rbpms*, *Pou4f1*, *Pou4f2*, and *Pou4f3* were identified as RGCs and separated from the other classes for further analysis. RGCs were then annotated with adult type labels via a supervised classification approach using gradient boosted trees (XGBoost) (Chen and Guestrin, 2016). In this approach, a classifier learns the transcriptomic signatures of cell types from a reference dataset (e.g. P56 cell types) to then assign type labels to cells in a test dataset (e.g. P7 cells). Details of the implementation can be found in (Shekhar et al., 2022). Briefly, two classifiers are trained on the reference (from which labels are learned) and test (to which labels are applied) datasets independently to identify features that contribute the highest average information gain across decision trees. Then, the top 500 discriminatory features are used in the training of a third classifier on the reference dataset. This classifier is then used to assign labels to cells in the test dataset. The classifier assigns a vector of probabilities to each cell, where entry *p_i_* of the vector corresponds with the probability that the cell belongs to label *i*. The label *i* that maximizes *p_i_* is then chosen as the label for the cell, unless the maximal *p_i_* is less than a threshold of 0.05. The same label transfer method was used to assign P56 types to P5 cells. XGBoost was run using a multi-log loss function and hyperparameter values `eta = 0.6`, `max_depth = 6`, and `subsample = 0.6`.

### Analysis of wave-dependent changes in gene expression

For all genes included in the four datasets, log2-fold changes were calculated for each pair of datasets. In the assessment of global gene expression changes, expression values were averaged over all cells in each dataset. For a gene *g* with average normalized expression *x_gi_* at Condition *i*, the log2-fold change between conditions *a* and *b* was calculated as 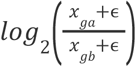 where ɛ = 0. 01 is a small pseudocount used to avoid undefined or extreme values. The same calculation was done to find inter-replicate log2-fold changes.

To assess type-specific gene expression changes, a similar calculation was done with average gene expression values representing type-condition pair averages rather than condition averages. Additionally, a larger pseudocount was used to account for the smaller quantity of cells within each type. That is, for a gene *g* with average normalized expression *x_gij_* at condition *i* in type *j*, the log2-fold change between conditions *a* and *b* in type *t* was calculated as 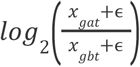

To investigate wave-dependent, type-specific changes in expression, we focused on changes between P7 WT and β2KO. For each type, a differential expression test (Wilcoxon rank-sum) was performed on the set of genes with a log2-fold change of at least 1.2 in magnitude. Genes with an adjusted p-value of at most 0.05 were then selected for Gene Ontology (GO) analysis.

Statistical overrepresentation analysis was performed using the PANTHER web interface (Ashburner et al., 2000; Thomas et al., 2022; The Gene Ontology Consortium et al., 2023) to identify GO terms associated with each set of wave-regulated genes. The test utilizes an input test set and background set to assign a p-value to each GO term. For a given GO term, let *k* represent the number of genes in the test set belonging to the GO term, *n* the length of the full test set, *K* the number of genes in the background set belonging to the GO term, and *N* the length of the full background set. Then, the probability that the term is enriched in the test set is calculated as

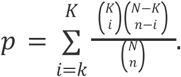

Two tests were performed to assess the GO terms enriched in both the β2KO upregulated and downregulated gene sets. For each of these tests, the test set was the set of genes that was found to be significantly upregulated or downregulated in P7 β2KO compared to P7 WT, and the input background set was the set of genes that passed the log2-fold change threshold. From this test, we identified those GO terms with adjusted p-values less than 0.1 and selected a few that are relevant to neuronal circuit development to highlight.

### Analysis of bulk RNAseq data

Bulk RNAseq libraries for three WT and four β2KO samples were aligned to UCSC mm10 genome assembly using STAR v2.7.11b (Dobin et al., 2013), and mapped reads were counted using featureCounts (Liao et al., 2014). Differential expression analysis was then performed using the PyDESeq2, a python implementation of DESeq2 (Love et al., 2014; Muzellec et al., 2023). Briefly, the count matrix was filtered to include only genes for which at least 10 reads were detected. Count matrices were then normalized using the median-of-ratios method of DESeq2. To model read counts with a negative binomial distribution, gene-level dispersions were calculated using the normalized matrix. A curve was fit across all dispersion values as a function of mean normalized counts, which was used to shrink dispersion values toward their predicted values and obtain the final dispersion parameters to be used in the model. Log2FCs changes were then computed between the WT and β2KO conditions, and these values were shrunk in a similar manner to the dispersion values to manage the high variance of log2FCs for genes with low read counts. Finally, Wald tests were performed on the log2FC estimates to determine significance of expression changes and identify differentially expressed genes. Genes were labeled as significantly differentially expressed if they had log2FC > 1 and p_adj_ < 0.05.

## Results

### β2KO exhibits normal RGC diversification

To assess the impact of cholinergic retinal waves on the RGC transcriptome, we used droplet-based scRNAseq (10X Chromium v3) to profile retinas from WT and β2KO mice at P7 (**Fig. 1A**). We chose P7 as it represents the final phase of wave reduction and precedes the onset of premature glutamatergic waves in β2KO (Bansal et al., 2000). As RGCs comprise fewer than 1% of all retinal cells, we enriched them using CD90-conjugated magnetic beads, following previously established protocols (**Methods**; Shekhar et al., 2022). We obtained 47,361 transcriptomes passing quality metrics, which includes the number of detected genes per cell (>700) and transcripts per cell (>5,000) detected, and a low proportion of mitochondrial transcripts per cell (< 10%). We classified these cells into major retinal classes based on the selective expression of marker genes (**Fig. 1B**). RGCs could be identified based on their high expression of the gene *Slc17a6*, which encodes the glutamate transporter VGLUT2; *Rbpms*, which encodes an RNA-binding protein; and the transcription factor encoding genes *Pou4f1-3* (**Fig. 1B**). Expression levels of *Chrnb2,* the gene encoding the β2 subunit of nicotinic acetylcholine receptor, were significantly lower in β2KO RGCs compared to WT RGCs, but were not zero (**Fig. 1B**). A closer examination revealed that the low basal level of expression in β2KO was due to reads mapping to another gene, *Adar*, whose 3’ end overlaps with the 3’ end of *Chrnb2* (**Fig. 1-1**). The remaining cell classes comprised amacrine cells, bipolar cells, Müller glia, endothelial cells, and microglia (**Fig. 1C**). Overall, ∼99% of cells in the WT dataset (n=27,274 cells) and ∼93% in the β2KO dataset (n=16,684 cells) were identified as high-quality RGCs (>1800 genes/cell and >5,000 transcripts/cell), indicating a successful enrichment. The rest of our analyses focuses exclusively on RGCs.

**Figure 1.**
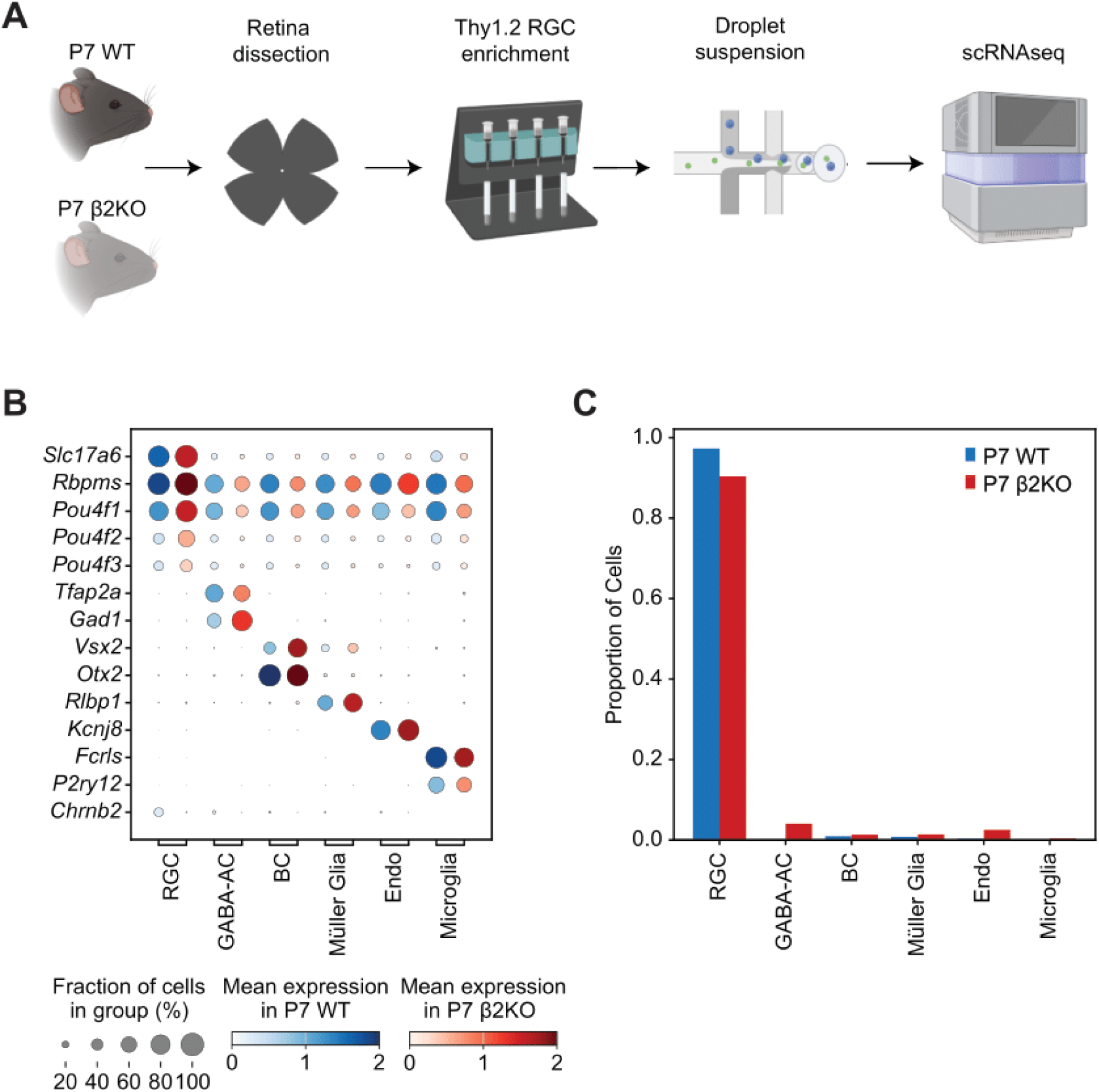
Study overview and enrichment of RGCs. **(A)** Overview of RGC enrichment and scRNAseq profiling from WT and β2KO mice. **(B)** Dotplot showing expression patterns of well-known markers (rows) in retinal cell classes (columns) within either condition. For each cell class, the expression levels in the WT and β2KO correspond to blue and red dots, respectively. The size of each dot corresponds to the percentage of cells in each class expressing the marker, and the color indicates the normalized expression level (see legend, bottom). **(C)** Barplot showing the relative proportion of major retinal cell groups (x-axis) in either dataset. BC, bipolar cells; GABA-AC, GABAergic amacrine cells; Endo, Endothelial cells. Note that we did not detect appreciable numbers of photoreceptors and horizontal cells, two prominent retinal cell classes.

To begin, we applied standard dimensionality reduction and clustering approaches to identify transcriptomically distinct clusters of RGCs in both WT and β2KO samples (**Methods; Fig. 2-1**). To relate transcriptomic clusters to RGC types defined previously, we trained a supervised classifier on a published atlas of adult (P56) RGCs (Tran et al., 2019) and applied this classifier to each P7 RGC (**Figs. 2A, B; Methods**). Based on an ensemble voting procedure, the classifier assigns each P7 RGC a P56-type label (C1-C45) and an associated measure of confidence. Using this procedure, most P7 RGCs (>91 % in WT and >96% in β2KO) could be mapped to an adult type with high confidence (**Methods**). Cells mapping to the same RGC type also co-clustered in the reduced dimensional space (**Fig. 2B**). Because several molecularly defined types have been mapped onto morphologically and functionally defined types in recent studies (Tran et al., 2019; Goetz et al., 2022; Huang et al., 2022), we were able to identify cells corresponding to well-known groups of types, such as alpha (α) RGCs (Krieger et al., 2017), ON-OFF and ON direction-selective RGCs (DSGCs), intrinsically photosensitive (ip) RGCs, and FoxP2 RGCs (Rousso et al., 2016). The P7 datasets contained all 45 molecularly defined RGC types.

**Figure 2.**
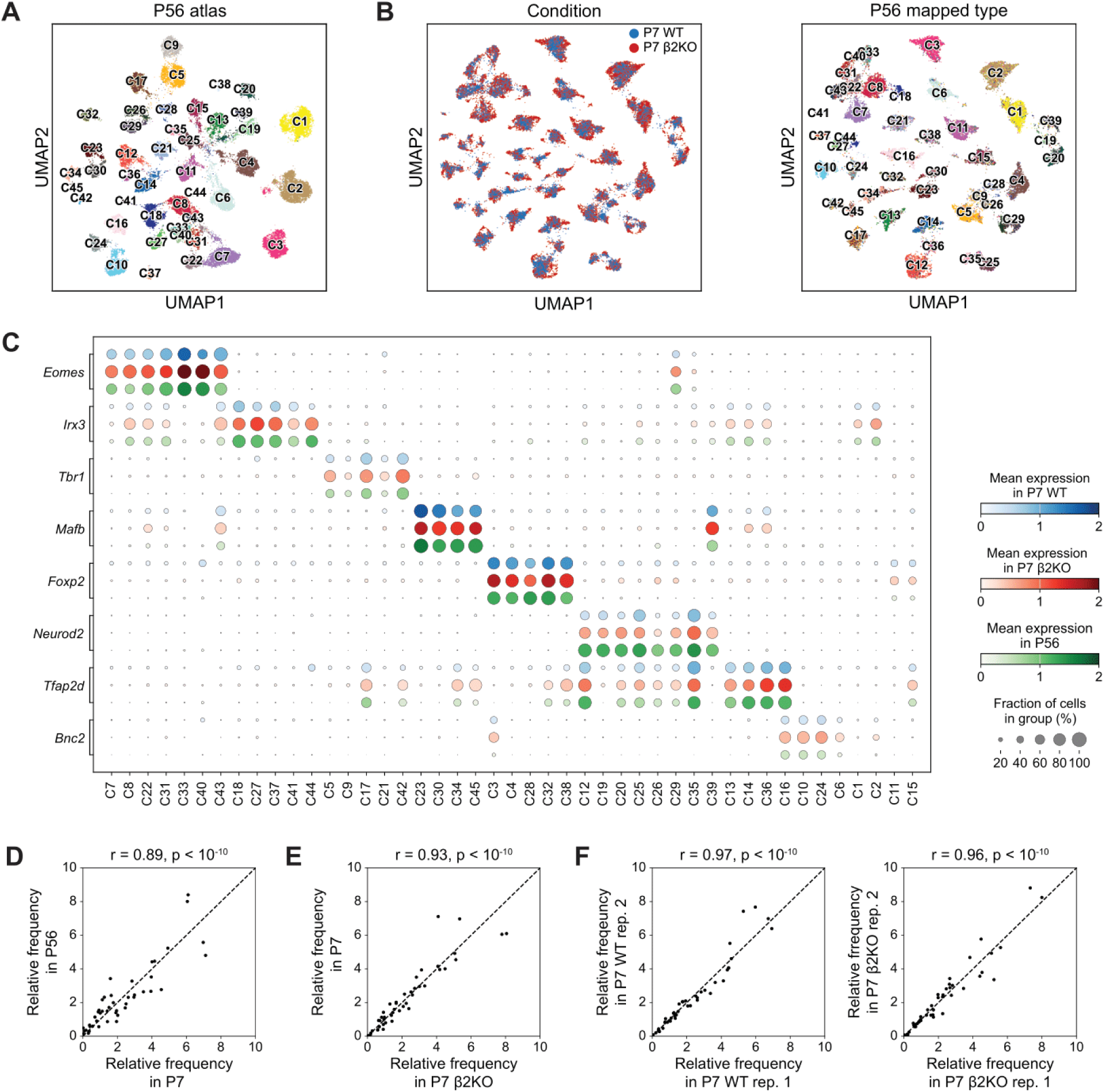
Disruption of cholinergic waves minimally impacts RGC diversification. **(A)** Uniform Manifold Approximation and Projection (UMAP) embedding of adult (P56) RGCs from (Tran et al., 2019) colored by molecularly defined types C1-C45. **Table 1-1** relates type IDs to commonly used type names. **(B)** Integrated UMAPs of cells from WT and β2KO RGCs at P7, colored by condition (left) and inferred type label (right) using a supervised classifier trained on P56 data. **(C)** Dotplot showing the expression patterns of subclass-specific transcription factors (row blocks) in each RGC type (columns). Within each row block, the three rows correspond to P7 WT (blue), P7 β2KO (red), and P56 WT (green). **(D)** Scatter plot comparing the relative frequency, *i.e.* cells in a cluster, between P56 WT and P7 WT. Each dot represents one RGC type. **(E)** Same as D, for P7 WT and P7 β2KO. **(F)** Same as D, for two replicates of P7 WT (left) and two replicates of P7 β2KO (right).

We verified these classifications *post hoc* by comparing cell type-specific gene expression patterns across the three datasets: P7 WT, P7 β2KO, and P56 WT. In previous works, we have shown that the 45 RGC types can be grouped into eight “subclasses” based on the selective expression of transcription factor (TF) encoding genes (Shekhar et al., 2022; Whitney et al., 2023). The three datasets showed consistent expression patterns of these subclass-defining TFs (**Fig. 2C**), as well as of other TFs and recognition molecules (**Fig. 3-1**). Although some types varied in composition between P7 WT and P7 β2KO, and between P7 WT and P56 WT, we found that the overall rank order of the relative frequency, which corresponds to the number of cells in a cluster, was highly correlated across the datasets (Pearson R > 0.8, p < 10^-15^) (**Fig. 2D-E**). Notably, the variations in type-frequencies between P7 WT and P7 β2KO were comparable in magnitude to those observed between biological replicates within the same condition (**Fig. 2F**), suggesting that these differences may not be biologically relevant. Taken together, the presence of all types in WT and β2KO, and the absence of any apparent differences in frequency composition suggest that retinal waves do not impact the diversification of RGCs. However, the disruption of retinal waves may still lead to altered gene expression without affecting the differentiation of types, which we now explore.

### Wave-dependent transcriptomic changes are modest globally but significant at the type-specific level

To assess the magnitude of changes due to wave disruption, we first computed differentially expressed genes (DEGs) along the course of normal development by comparing WT RGCs at P7 with WT RGCs at P56 and at P5. The P5 data was obtained from our previous study (Shekhar et al., 2022) using an older droplet-based technology (Macosko et al., 2015). P5 and P7 lie in the period over which cholinergic retinal waves occur. RGCs elaborate dendrites and make synapses with bipolar cells and amacrine cells in the inner plexiform layer before the detection of the first light responses (∼P10). The retina is mature by P56. Integration of the published P5 and P56 RGC datasets with P7 WT and P7 β2KO datasets revealed that cells mixed well and separated by their adult type labels, even though the type labels were not used explicitly to guide the integrated clustering procedure (**Fig. 3A**).

**Figure 3.**
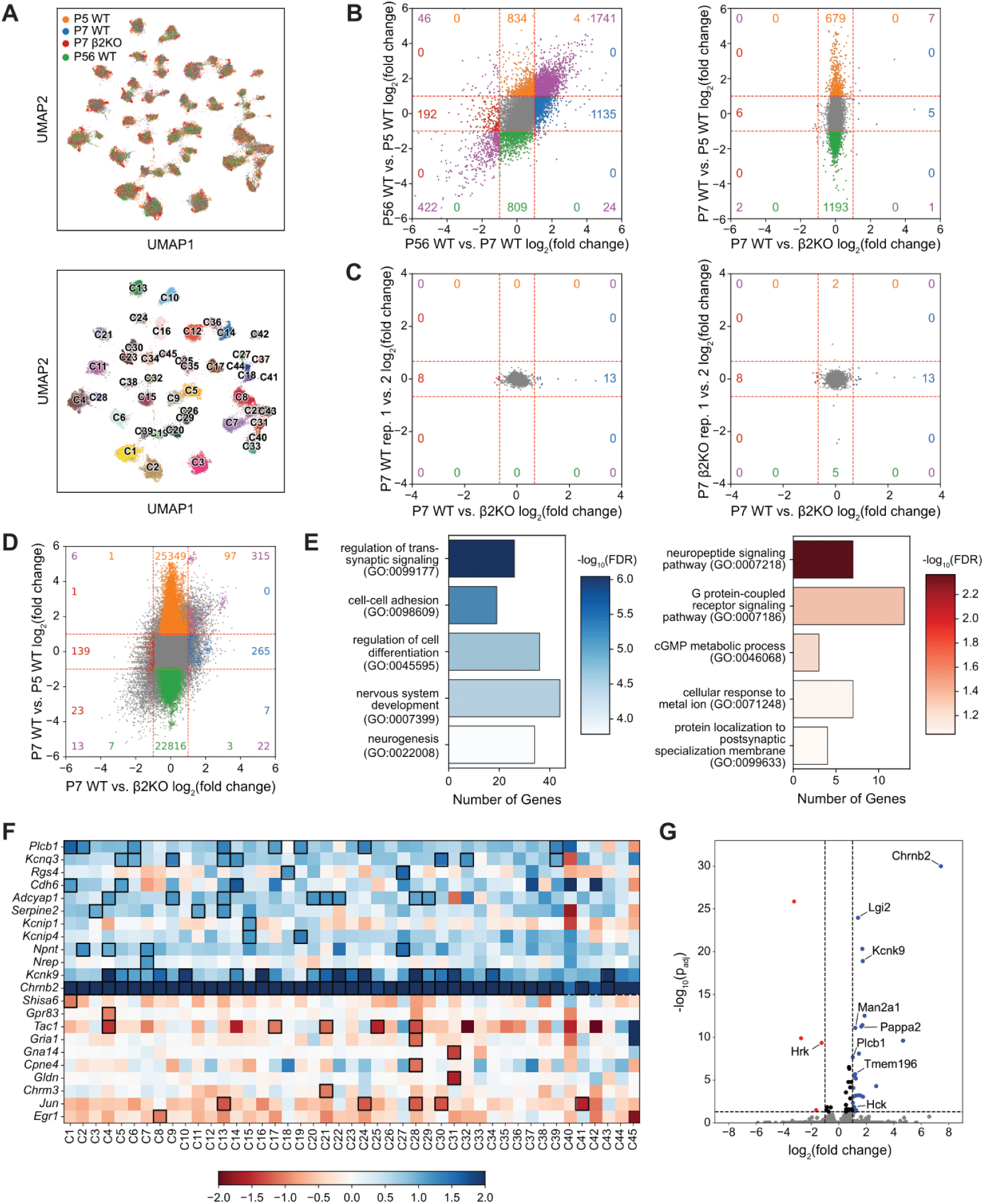
Global and type-specific wave-dependent transcriptomic changes. **(A)** Integrated UMAP of cells from P5 WT, P7 WT, P7 β2KO, and P56 WT colored by the dataset of origin (top) and mapped adult type (bottom). **(B)** Comparisons of log2-fold changes (log2FC) in global gene expression across datasets. Left panel compares WT datasets: P56 vs. P5 (y-axis) with P56 vs. P7 (x-axis). Right panel compares P7 WT vs. P5 WT (y-axis) with P7 WT vs. P7 β2KO (x-axis). Colored dots represent significantly up/downregulated genes (log2FC > 1, Wilcoxon rank-sum test adjusted p-value < 0.05). Each color represents significant up/downregulation in one comparison, and purple represents genes that are significantly up/downregulated across both comparisons. For comparisons W vs. X on the x-axis and Y vs. Z on the y-axis, genes upregulated in W vs. X are blue, genes downregulated in W vs. X are red, genes upregulated in Y vs. Z are orange, genes downregulated in Y vs. Z are green, and genes that fall into any two of these categories are purple. Colored numbers represent the count of genes falling in each category. **(C)** Comparisons of log2FC in global gene expression between the two P7 conditions and between biological replicates. Left panel compares P7 WT replicate 1 vs. replicate 2 (y-axis) with P7 WT vs. P7 β2KO (x-axis). Right panel compares P7 β2KO replicate 1 vs. replicate 2 (y-axis) with P7 WT vs. P7 β2KO (x-axis). Colors as in (B). **(D)** Comparisons of log2FC in type-specific gene expression between P7 WT vs. P5 WT (y-axis) and P56 vs. P5 (y-axis) with P56 vs. P5 (x-axis). Each dot corresponds to a gene-type combination (e.g. *Chrnb2* in C33). Colors as in (B). **(E)** GO terms overrepresented in the genes significantly downregulated (left, blue) and upregulated (right, red) in β2KO. **(F)** WT vs. β2KO log2FC changes in expression of example genes contributing to the GO terms shown in E. Black boxes indicate significant up/downregulation of a gene in a type (log2FC > 1, Wilcoxon rank-sum test p_adj_ < 0.05). **(G)** Volcano plot depicting differentially expressed genes between WT and β2KO identified using bulk RNAseq. Genes found to be significantly differentially expressed in at least one type using the scRNAseq data are labeled (a full list of genes is given in Fig. 5-1). For both scRNAseq and bulk RNAseq, each sample/replicate corresponds to RGCs pooled from different mice, making each one a biological replicate.

We first assessed global transcriptomic changes (*i.e.*, without regard to cell type identity) across each pair of datasets, beginning with developmental changes. At a threshold of log2-fold change with log2FC > 1 and adjusted p-value p_adj_ < 0.05 (Wilcoxon rank-sum test), there were 3880 differentially expressed genes (DEGs) between P5 and P56 WT and 3560 DEGs between P7 and P56 WT. 2233 genes were common among the two sets with a statistically significant overlap (hypergeometric test, p < 10^-10^), highlighting that, as expected, P5 and P7 are more similar to each other than each of them is to P56 (**Fig. 3B, left**). We also found 1882 DEGs between P5 and P7, suggesting considerable transcriptional regulation between these two temporally proximal time points (**Fig. 3B, right**). By contrast, there were only 21 genes that were globally differentially regulated between P7 WT and P7 β2KO. This wide disparity implies that wave-dependent changes are modest even when compared to changes that occur during two days of normal development, as evidenced by the greater spread along the y-axis in the right scatterplot of **Fig. 3B**. On the other hand, we identified no DEGs between biological replicates within P7 WT or P7 β2KO, suggesting that these modest changes are likely biologically meaningful (**Fig. 3C**).

We hypothesized that the disruption of cholinergic waves may impact RGC types non-uniformly, such that an impacted gene is dysregulated in only a few RGC types. Combined with several unaffected types, these changes may be washed out in global comparisons. Accordingly, within each type, we computed DEGs that were wave-dependent or developmentally regulated. Though developmental changes still dominate, we identified many type-specific, wave-dependent DEGs that did not appear in the global comparisons (**Fig. 3D**). Overall, we found 238 genes that were significantly upregulated or downregulated (log2FC > 1, p_adj_< 0.05) in individual types. On average, each gene was regulated in 3-8 types, and each type had 18 ± 9 DEGs. To understand what biological processes are represented amongst these type-specific, wave-dependent genes, we performed a Gene Ontology (GO) enrichment analysis on the DEGs (Ashburner et al., 2000; Thomas et al., 2022; The Gene Ontology Consortium et al., 2023). The top terms enriched among both the upregulated and downregulated genes included processes related to cell-cell adhesion, and synaptic and neuropeptide signaling, indicating that type-specific changes in gene expression likely have differential developmental impact on different visual feature detector pathways (**Fig. 3E**). Contributing to the enrichment of these processes were several genes that encode potassium channels and subunits of neurotransmitter (e.g. GABA, glutamate) receptors, which are highlighted in **Fig. 3F** and **Fig. 4-1**.

To validate our findings and ensure that these modest transcriptomic changes were not an effect of the high dropout rate of scRNAseq, we performed bulk RNAseq on RGCs from WT and β2KO mice (**Methods**). A differential expression analysis of this data again revealed modest gene expression changes between the two conditions, with only 32 genes appearing as significantly up/downregulated between the WT and β2KO (**Figs. 3G, 5-1**). Among these genes, 9 (hypergeometric test, p < 10^-10^) were also identified as significantly up/downregulated in at least one type in the single-cell analysis, providing independent support that these genes are indeed regulated by waves.

### Transcriptome of horizontal motion-preferring direction-selective ganglion cells minimally changed in β2KO

Our previous work has shown that the β2KO has a dramatic reduction in direction-selective responses of horizontal-motion preferring DSGCs (Tiriac et al., 2022). Mouse DSGCs can be classified into two main subclasses, ON-OFF and ON, via their mode of light responses. ON-OFF DSGCs can be further divided into four types (dorsal (D), ventral (V), nasal (N), and temporal (T)) and ON DSGCs into three (D, V, and N) based on their directional preference (Oyster and Barlow, 1967; Sun et al., 2006; Elstrott et al., 2008; Yonehara et al., 2009; Dhande et al., 2013). In the β2KO, we saw a reduction in the direction selective responses of N- and T-ON-OFF DSGCs as well as N-ON DSGCs, while all vertical DSGCs retained their tuning. Importantly, we should note that in the β2KO, N-ON-OFF DSGCs still existed, but had only lost their tuning properties.

To assess the differential impact of wave disruption on the different DSGC types, we first had to identify the cell types transcriptomically at P7. At P56, three clusters of ON-OFF DSGCs have been previously reported: C12 as N-ON-OFF DSGCs, C16 as a mix of D- and V-ON-OFF DSGCs, and recent studies identifying C24 as T-ON-OFF DSGCs (Goetz et al., 2022; Huang et al., 2022). Through supervised analysis, C16 can be subclassified into dorsal and ventral types based on expression of *Calb1* and *Calb2* (D: high *Calb1*/low *Calb2*, V: high *Calb2*/low *Calb1*). C10 in the adult atlas has recently been mapped to ON-DSGCs in both mouse (Goetz et al., 2022; Huang et al., 2022; Toma et al., 2024), and primate (Hahn et al., 2023; Wang et al., 2023), but further subclustering of C10 into the D, V and N varieties was unsuccessful in the adult data (**Fig. 4A**).

**Figure 4.**
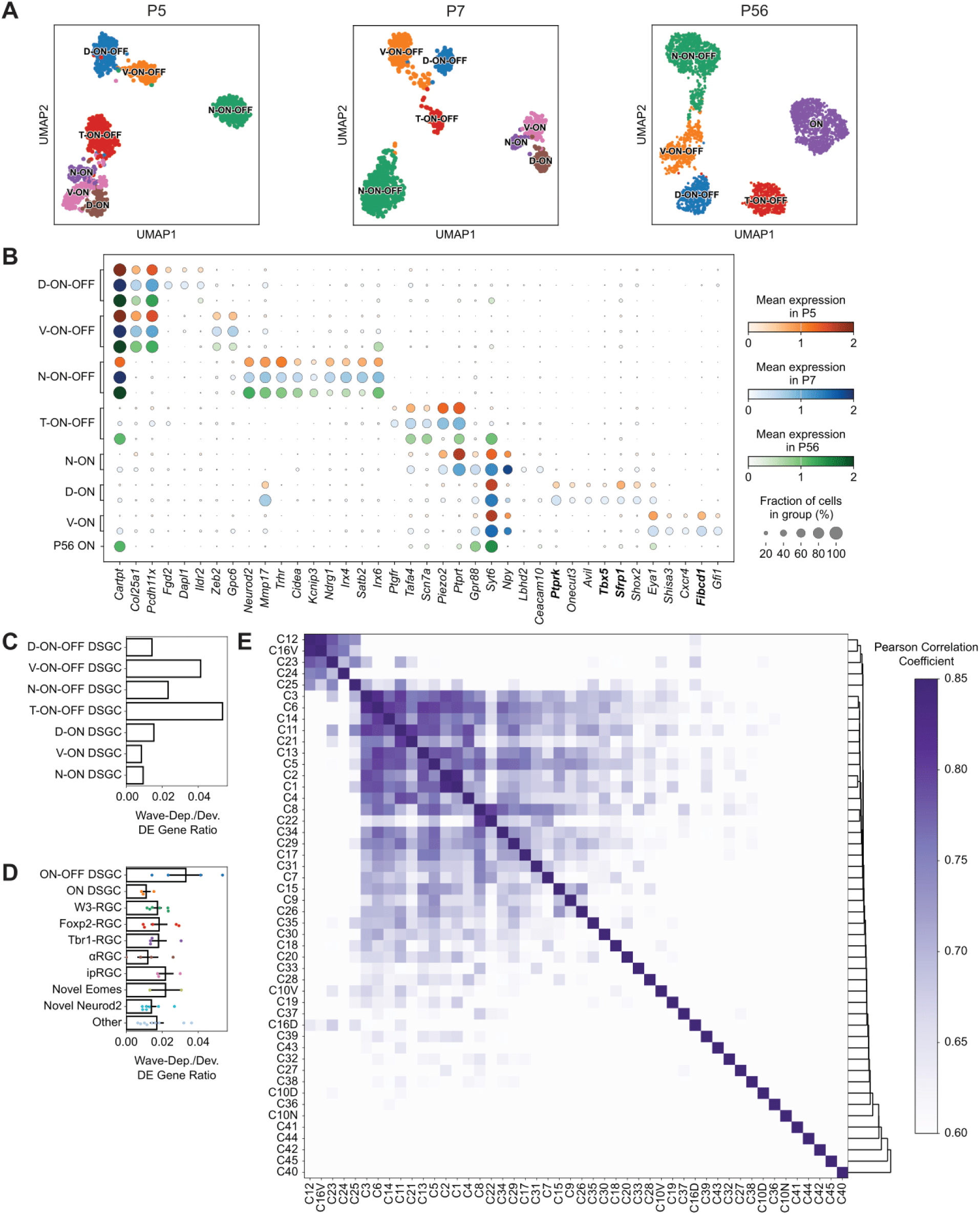
Classification of DSGCs and analysis of subclass-specific, wave-dependent regulation. **(A)** UMAP representations of DSGCs at P5, P7, and P56, colored by DSGC type. **(B)** Expression of marker genes for each DSGC type across the age conditions. Only P5 and P7 are represented for individual ON-DSGC types, with the bottom row showing expression in the unclustered ON-DSGC group at P56. Genes in bold correspond to markers for ON-DSGCs during development. **(C)** Ratios of wave-dependent (P7 WT vs. P7 β2KO) to developmental (P7 vs. P5) DE gene counts per DSGC type. **(D)** Ratios of wave-dependent (P7 WT vs. P7 β2KO) to developmental (P7 vs. P5) DE gene counts per RGC subclass, with points representing individual types. Bars lengths represent mean ± SEM. **(E)** Pairwise correlations in gene log2FC between RGC types, ordered based on hierarchical clustering. Darker colors represent stronger correlations. **Table 1-1** relates cluster IDs (Tran et al., 2019) to commonly used type names. DSGC clusters split here (C16 and C10) are appended with the letter corresponding to their preferred direction (e.g. C16V represents V-ON-OFF DSGCs).

By contrast, the D, V, and N types among ON DSGCs could be molecularly distinguished at earlier stages (**Fig. 4A**). We found that immature RGCs mapping to C10 in the adult formed three clusters during development, which mapped 1:1 between P5 and P7. **Figure 4B** summarizes the molecular taxonomy of all DSGC types at the three ages. Moreover, these ON DSGC gene signatures at P5 and P7 were consistent with a recent study which analyzed molecular diversity of ON DSGCs by applying Smart-seq2 to FACS-enriched RGCs obtained from a knockin transgenic mouse line that selectively labels ON DSGCs (Al-Khindi et al., 2022). Finally, results suggest that while ON-OFF marker distinctions persist from P5 into adulthood, those associated with ON DSGCs are lost during maturation (**Fig. 4B**). As all our datasets employ droplet-based scRNAseq, these results also argue against the possibility that our inability to resolve ON DSGC diversity in adult mice is due to technical sampling limitations.

With the DSGC classifications in hand, we examined whether the transcriptome of horizontal motion-preferring DSGCs is more susceptible to waves than that of vertical ones. To quantify the magnitude of wave disruption, we use the ratio of wave-dependent DE gene counts to developmental DE gene counts as a metric for susceptibility to waves. A comparison of DSGC types reveals minor variations in this ratio from type to type, with these differences being very small in magnitude, suggesting that transcriptomically no DSGC type was more susceptible to wave disruption (**Fig. 4C**).

We then expanded this analysis to other RGC subclasses, including *Foxp2*-RGCs, ipRGCs, and αRGCs. We calculated wave-dependent/developmental DE count ratios for all RGC types, then grouped these ratios by subclass. While subtle differences were observed, there was no significant variation in mean ratios across subclasses (one-way ANOVA, p = 0.13) (**Fig. 4D**). Though the magnitude of wave-dependent changes does not seem to differ significantly across subclasses, it may be the case that waves regulate similar gene expression changes in certain RGC subclasses. To investigate this, we computed pairwise correlations in gene log2FCs across all RGC types (**Fig. 4E**). While there was a stronger correlation in wave-dependent expression changes between types within certain subclasses, namely in ON-OFF DSGCs and W3-RGCs, most RGC types had different molecular programs impacted due to wave disruption.

### Expression of a leak potassium channel is downregulated in β2KO

To assess the functional impact of wave-dependent gene regulation on RGCs, we focused on *Kcnk9*, a gene downregulated in several β2KO RGC types, namely αRGCs (**Fig 3F, 5A**). *Kcnk9* encodes TASK-3, a two-pore-domain leak potassium channel, which has been shown to contribute to neuronal excitability in αRGCs (Wen et al., 2022). *Kcnk9* mRNA expression was downregulated in all four types of αRGCs, though statistical significance could only be established for αON-Sustained RGCs due to sampling limitations (**Fig 6-1**). We used *in situ* hybridization (ISH) to validate the downregulation of *Kcnk9* in β2KO RGCs. Combined with immunohistochemistry (IHC) for SMI32, an αRGC marker, these experiments also showed a robust decrease in *Kcnk9* expression within SMI32^+^ αRGCs (**Fig 5B-D**).

**Figure 5.**
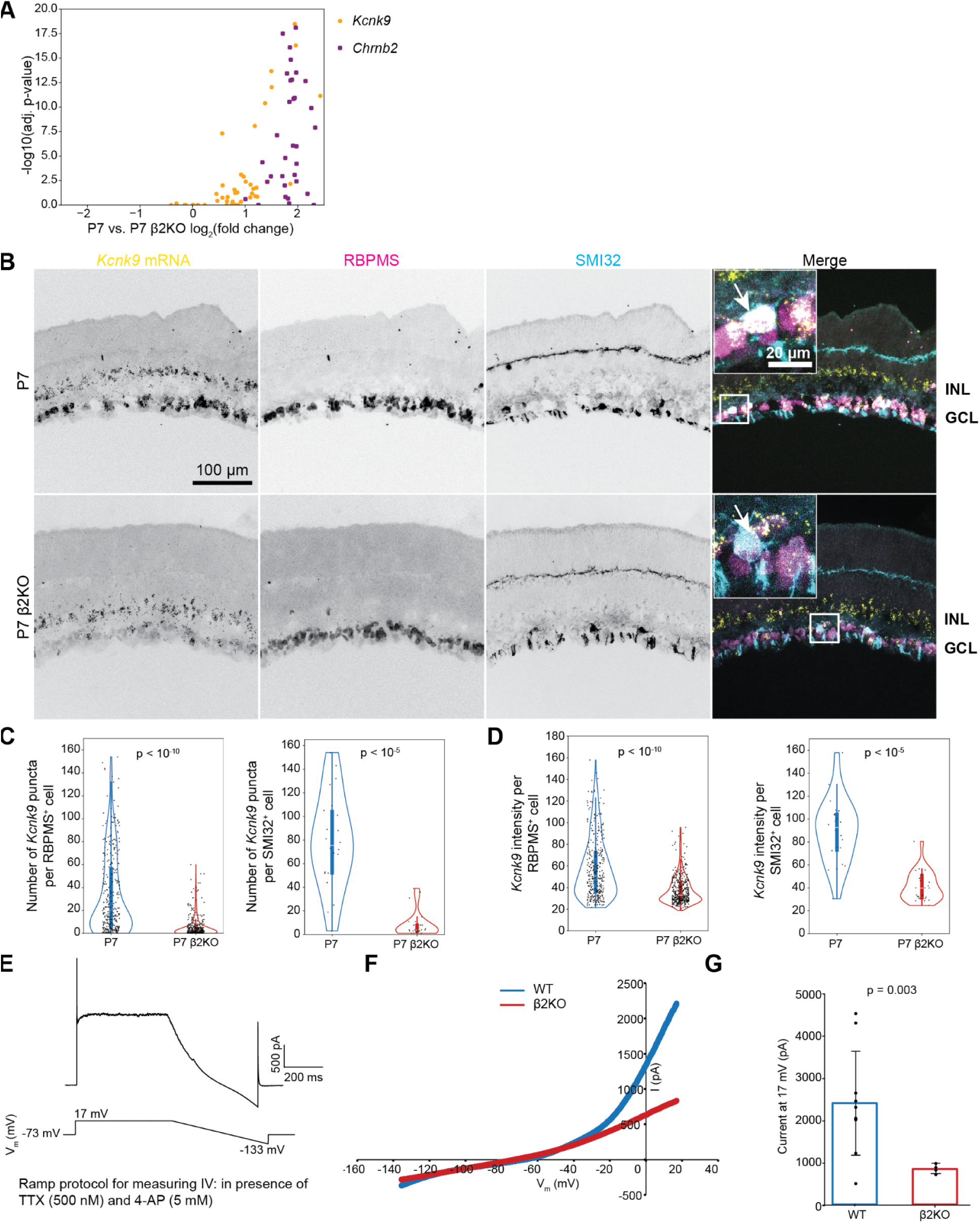
Decrease in the expression and function of leak channel TASK3 in β2KO alpha RGCs. **(A)** Volcano plot depicting type-level log2FCs and Wilcoxon rank-sum test adjusted p-values for *Chrnb2* (purple squares) and *Kcnk9* (orange circles). Each dot corresponds to an RGC type. **(B)** ISH for *Kcnk9* mRNA (yellow) on P7 WT and β2KO retinal slices. IHC staining to identify RGCs labeled with RBPMS (magenta) and αRGCs labeled with SMI32 (cyan). Inserts depict high-magnification images where arrows indicate high (in P7 WT) or low (P7 β2KO) expression of *Kcnk9* in RBPMS^+^SMI32^+^ αRGCs. **(C, D)** Quantification for number of *Kcnk9* puncta and fluorescence intensity in RBPMS^+^ cells (left) and SMI32^+^ cells (right) in P7 WT and β2KO. Each data point represents value from an individual cell. p < 10^-10^ for RBPMS^+^ cells and p < 10^-5^ for SMI32^+^ cells (Mann-Whitney U Test). **(E)** Voltage-clamp ramp protocol used previously to characterize TASK3 (Wen et al., 2022) (bottom) and example response trace from αRGC (top). **(F)** Example current-voltage curve of αRGCs from P7 WT (blue) and P8 β2KO (red) mice, P7-P12. **(G)** Summary data based on current recorded at Vhold = 17 mV in αRGCs WT (n=10) and β2KO (n=4), P7-P12. Each data point represents the value from an individual cell. Error bars = standard deviation, p value determined by Welch’s two-sample t-test.

To assess the impact on leak channel-associated conductance in αRGCs, we conducted voltage-clamp recordings. αRGCs were identified in a flat-mounted retina as RGCs with soma diameters of >15 μm. To isolate the leak channel-associated conductance, these recordings were done in the presence of tetrodotoxin (TTX, 500nM) to block voltage-gated sodium channels and 4-aminopyridine (4-AP, 5 mM) to block voltage-gated potassium channels (**Figs. 5E, F**). β2KO αRGCs showed significantly reduced leak channel-associated conductance compared to WT (**Fig. 5G**), consistent with reduced expression of TASK3. These data show that transcriptomic changes in *Kcnk9* expression in β2KO alter outwardly rectifying potassium currents and suggest that disrupting cholinergic retinal waves likely increases the intrinsic excitability of αRGCs, a potential form of homeostatic regulation.

## Discussion

By performing scRNAseq on developing RGCs, we discovered that the disruption of cholinergic retinal waves in β2KO mice led to a subtle RGC-type-specific regulation of expression of several genes. Notably, wave-dependent changes in the RGC transcriptome were more subtle than developmental changes, in sharp contrast to the dramatic changes observed in retinorecipient brain regions following wave disruption (discussed below). We also report that despite an absence of direction selectivity in horizontal motion-preferring DSGCs in β2KO mice, there was a minimal impact on their gene expression profiles. Our data suggests that cholinergic wave-dependent molecular changes in retinal circuits do not prominently manifest at the transcriptomic level in all RGC types, highlighting the need to explore post-transcriptional mechanisms.

### The absence of cholinergic waves does not impact RGC differentiation

The conventional view of neuronal differentiation into diverse cell types is that it occurs largely independent of neuronal activity (Greig et al., 2013). However, several studies have revealed a role of spontaneous activity in neuronal diversification within various areas of the central nervous system. For example, in the *Xenopus* spinal cord, spontaneous high-frequency calcium activity initiates a signaling cascade that drives unspecialized neurons to acquire a GABAergic identity (Borodinsky et al., 2004; Marek et al., 2010). Similarly, varied firing patterns of spontaneous activity among olfactory sensory neurons have been shown to induce the expression of unique axon-sorting molecules, conferring distinct identities to these neurons and enabling their axons to segregate into specific glomeruli within the mouse olfactory bulb (Yu et al., 2004; Serizawa et al., 2006; Mobley et al., 2010; Williams et al., 2011; Nakashima et al., 2019). Comparable roles for spontaneous activity have been observed in the cochlea, where activity during the pre-hearing phase regulates the specification of type I spiral ganglion neurons (Shrestha et al., 2018; Sun et al., 2018). Even in later stages of development, there are examples of sensory activity contributing to cell type specification. Olfaction-driven sensory activity is crucial for the differentiation of specific interneuron populations within the mouse olfactory bulb (Bastien-Dionne et al., 2010; Tepe et al., 2018), while visual experience during the critical period plays a pivotal role in the development and spatial patterning of glutamatergic cell types in the upper layers of the mouse primary visual cortex (Cheng et al., 2022).

In the retina, RGCs diversify gradually, with the full complement of 45 molecularly distinct types emerging only by the end of the first postnatal week (P5) (Shekhar et al., 2022). Notably, this diversification is unaffected by activity changes induced by visual deprivation (Whitney et al., 2023). Here, we extend these findings to show that even early postnatal spontaneous retinal activity, occurring prior to vision-driven activity, does not impact the diversification of RGC types. These results suggest that RGC diversification is likely guided by an intrinsic, activity-independent mechanism, aligning with the classical view of neuronal differentiation.

### Differential impact of retinal cholinergic waves across the visual system

Recent studies have examined the importance of retinal waves on the transcriptome of neurons in V1 and SC. For instance, conditional deletion of the β2 subunit of nAChRs in Pax6α^+^ cells—specifically expressed in peripheral retina, leading to disruption of waves in this region, caused dramatic alterations in the developmental transcriptome of both excitatory and inhibitory cells and disrupted synaptic inputs onto these neurons in posterior V1 (Burbridge et al., 2024). Notably, they observed that pyramidal cells and Layer 1 interneurons had significant changes in a large number of genes, while other inhibitory neurons had modest changes. In a different study, pharmacological blockade of perinatal retinal waves using gap junction blocker carbenoxolone led to significant changes in over 1000 genes in neurons of the superior and deep layers of the SC (Guillamón-Vivancos et al., 2022). In contrast, our manipulation of spontaneous retinal activity using β2KO led to significant transcriptomic changes in only a modest number of genes, with each gene regulated in a small number of types. All RGC types exhibited differentially expressed genes, and no single type was more susceptible to wave disruption. These findings indicate that spontaneous retinal activity has a more pronounced transcriptomic impact on postsynaptic cells in downstream brain visual regions as compared to RGCs.

Our results contrast to a recent study in which overexpression of the potassium channel Kir2.1 in RGCs, leading to lowered frequency of cholinergic waves, showed alteration of the expression of a large number of genes (∼1500 genes) in RGCs at P8 using bulk RNAseq (Negueruela et al., 2024). While it may seem that these differences stem from the higher sensitivity of bulk RNAseq to detect subtle, graded changes in gene expression, the bulk RNAseq analysis performed in this study showed results that were consistent with our scRNAseq data. Thus the sequencing technology alone cannot account for the discrepancy. We propose that these differences are more likely due to the distinct strategies used to manipulate RGC activity or due to the enrichment method for RGCs prior to sequencing. Overexpression of Kir2.1 may have induced physiological changes due to the introduction of a large potassium conductance, resulting in more extensive transcriptomic alterations than those caused by the absence of cholinergic signaling in β2KO mice. Furthermore, the RGC enrichment strategy using Pou4f2-cre line might have also isolated late-stage RGC progenitors or included subset of cells from the inner nuclear layer (Simmons et al., 2016). Our approach mitigated potential contaminant cells using antibodies against CD90, which extracts RGCs at >90% purity, as shown in **Figure 1C** and several other studies (Liu et al., 2018; Rheaume et al., 2018; Peng et al., 2019; Tran et al., 2019; Shekhar et al., 2022; Whitney et al., 2023). These differences highlight how transcriptomic readouts can be influenced by both the method of activity manipulation and cell enrichment.

Though the impact of cholinergic waves on the overall RGC transcriptome we observed was limited, many of the dysregulated genes identified in our study are involved in developmental processes, indicating that retinal waves may have specific, yet unexplored roles in regulating critical mechanisms in different RGC types. For example, we observed that in certain RGC types, waves regulate the expression of *Kcnk9*, which encodes the leak potassium channel TASK3. This finding is particularly significant because *Kcnk9* was uniquely disrupted in β2KO RGCs, in contrast to other retinal activity disruption models where either the wave frequency is altered or mice are deprived of visual experience (Whitney et al., 2023; Negueruela et al., 2024). The role of TASK3 in modulating intrinsic excitability suggests an intriguing connection between the presence of retinal waves and the intrinsic excitability of RGCs.

### Disruption of cholinergic waves leads to modest transcriptomic changes across DSGC subtypes

Previously, we showed that disrupting cholinergic retinal waves impairs horizontal direction selectivity in the retina but preserves vertical-motion preference of DSGCs (Tiriac et al., 2022). This aligns with other studies showing that retinal waves specifically influence the horizontal optokinetic reflex (Wang et al., 2009). These results suggest that horizontal and vertical motion-preferring direction-selective circuits in the retina develop through distinct signaling pathways, some of which are regulated by activity.

The emergence of direction-selective circuits coincides with robust wave activity. The direction preference of DSGCs is based on their wiring with SACs. Presynaptic SAC processes provide greater synaptic inhibition to postsynaptic DSGCs during null direction motion. This is a result of two features: 1) individual processes of SACs are more strongly depolarized by outward motion and 2) more synapse formation between SAC processes that are oriented parallel to the null direction of DSGCs (Briggman et al., 2011; Mauss et al., 2017). Paired recording between SACs and DSGCs has revealed that the asymmetric inhibitory synaptogenesis occurs between P9 to P11 in the mouse retina (Wei and Feller, 2011; Yonehara et al., 2011; Morrie and Feller, 2015; Tworig et al., 2024). During this period, vertical- and horizontal-motion preferring DSGCs exhibit distinct transcriptome with numerous DEGs shown to be critical for synapse development in other contexts (Tworig et al., 2024).

In this study, we investigated whether any of the differentially expressed genes in DSGCs are regulated by neural activity and whether the transcriptomes of horizontal-motion preferring DSGCs are particularly susceptible to wave disruption. Conducting this analysis during development provided an advantage, as unlike in adults, distinct ON-DSGC subtypes form separate transcriptomic clusters in the developing retina (**Fig 4B**). Surprisingly, intersectional analysis revealed that none of these DEGs were wave regulated (**Fig. 4C**), suggesting that spontaneous activity-mediated DSGC wiring does not occur by modulating cell-intrinsic genetic programs. While this result may also reflect the limited resolution of single-cell sequencing, which captures only about 15% of the cell’s transcriptome (10X Genomics, n.d.), it is also possible that retinal waves exerts their influence through post-transcriptional or translational processes in DSGCs (West and Greenberg, 2011). Such activity-dependent mechanisms have been observed in the hippocampus, where activity regulates the local translation of matrix metalloproteinase-9, a protein crucial for regulating spine morphology in neurons (Dziembowska et al., 2012). Similar activity-dependent local translation may act in DSGC synapses, potentially influencing their wiring with SACs. The development of newer technologies, such as isolating specific synapses using mouse transgenic lines or performing proximity protein labeling followed by mass spectrometry, could help investigate this possibility (Shuster et al., 2022; Van Oostrum et al., 2023).

## Acknowledgments and funding sources

Some images were made using BioRender. We thank Dr. Justin Choi from the QB3/Functional Genomics Lab for assistance with scRNAseq and bulk RNAseq, RRID:SCR_022170. Funding sources include NIH EY028625 (KS), Society of Hellman Fellows (KS), McKnight foundation (KS), Glaucoma Research Foundation (KS), Weill Neurohub (RDS), NIH EY013528 (MF), and NIH EY019498 (MF).

## Extended Data

**Figure S1-1.**
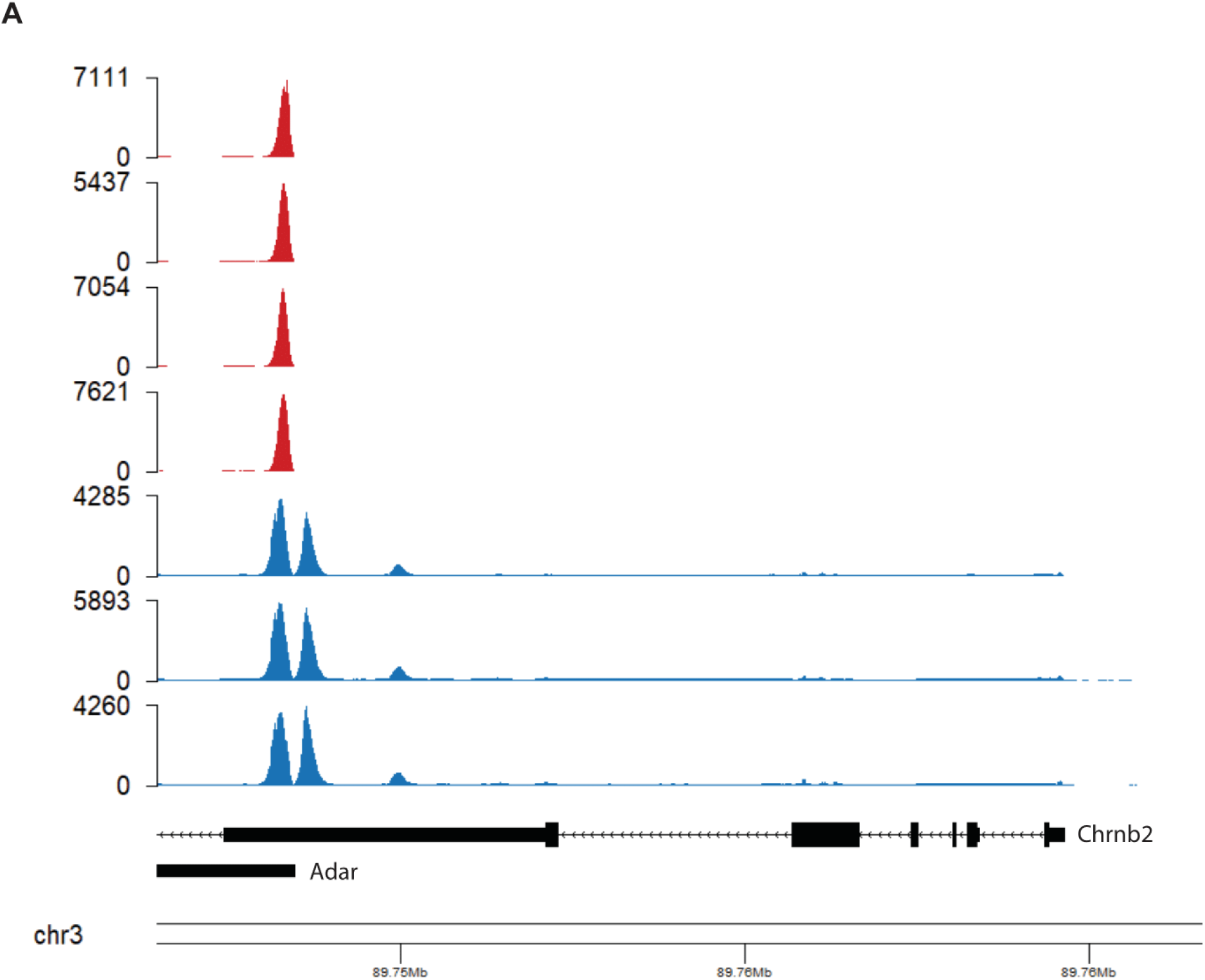
Reads from each sample mapped to the genome. The three WT samples are shown in blue (bottom), and the four β2KO samples are shown in red (top). A negligible amount of reads from the β2KO map to *Chrnb2*, and expression of the gene is likely due to reads from *Adar* mapping onto *Chrnb2*.

**Figure S2-1.**
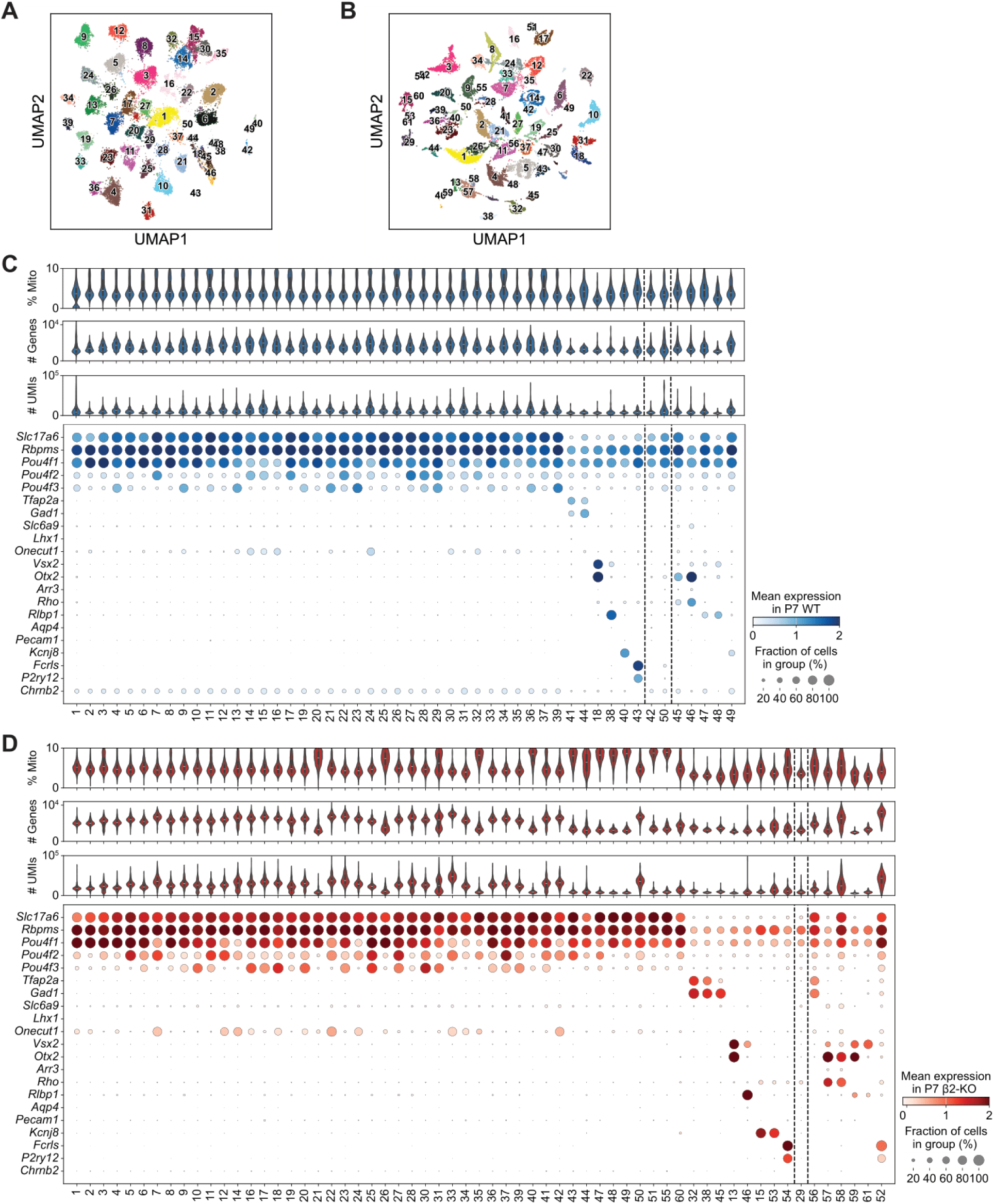
Classification and filtering of P7 cells. **(A-B)** UMAP representations of **(A)** P7 WT and **(B)** P7 β2KO cells colored by clusters assigned via the Leiden algorithm. **(C)** Cell quality metrics and expression profiles for P7 WT clusters. From top to bottom: mitochondrial proportion of genes per cell, number of genes expressed per cell, number of unique molecular identifiers (UMIs) per cell, and cluster expression of retinal cell class markers. Dotted lines separate clusters with clear class marker expression (left) from those expressing no retinal class markers (middle) and those expressing more than one class marker (right). **(D)** Same as (C) for P7 β2KO clusters.

**Figure S3-1.**
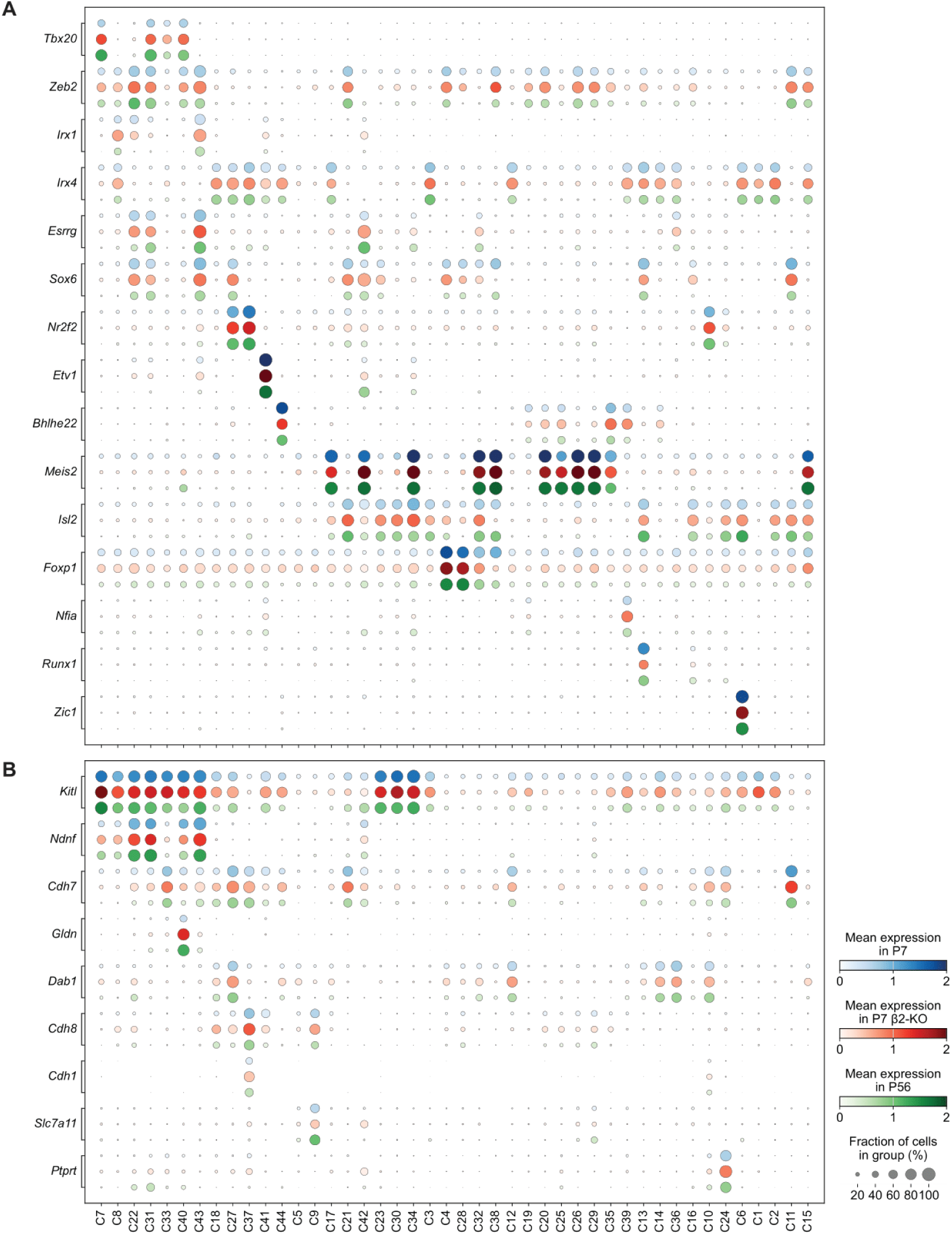
Congruent cell type-specific expression profiles of P7 WT, P7 β2KO, and P56 WT cells. Expression of (A) transcription factors and (B) cell adhesion molecules that are selectively expressed across RGC types.

**Figure S4-1.**
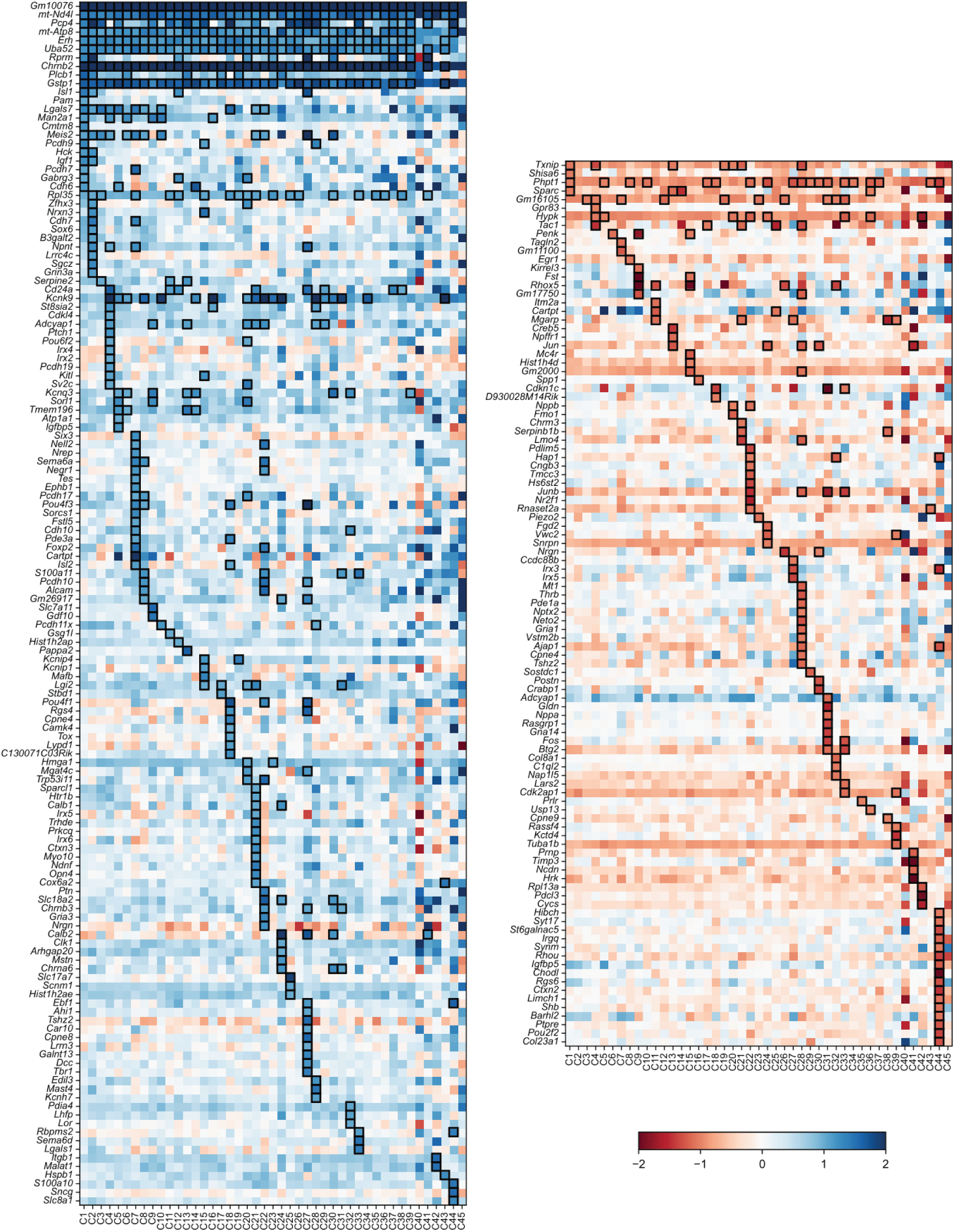
P7 WT vs. β2KO log2FC in all genes found to be significantly up/downregulated in at least one type. Black boxes indicate significant up/downregulation of a gene in a RGC type. (log2FC > 1, Wilcoxon rank-sum test adjusted p-value < 0.05). Blue corresponds to higher expression in WT and red corresponds to higher expression in β2KO.

**Figure S5-1.**
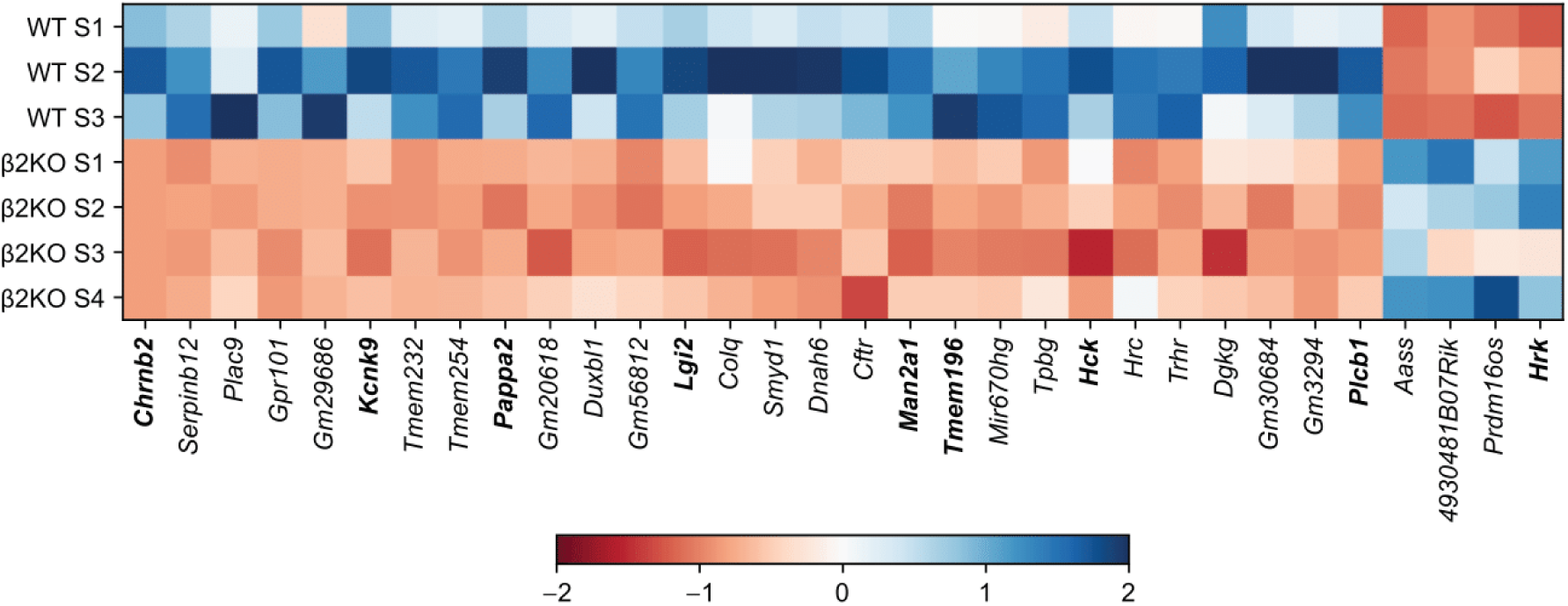
Normalized, z-scored expression of genes found to be differentially expressed using bulk RNAseq data. Genes identified as significantly differentially regulated in at least one type in the single-cell analysis are listed in bold.

**Figure S6-1.**
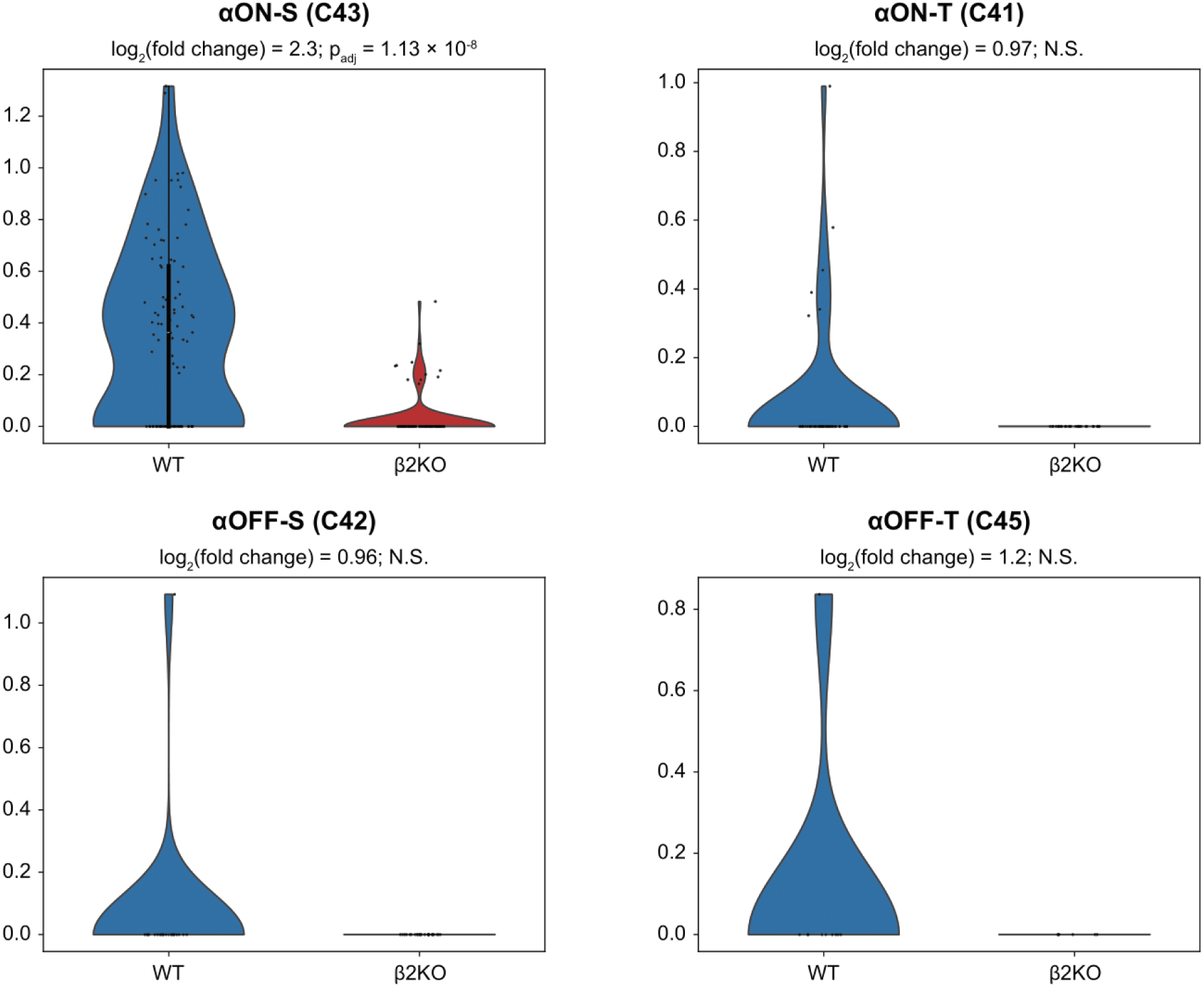
Downregulation of *Kcnk9* in αRGCs. Average expression of *Kcnk9* is higher in the WT condition for all αRGC types, but this difference is only found to be significant (Wilcoxon rank-sum test adjusted p-value < 0.05) in αON-S RGCs due to low sampling of other αRGCs. Note that fold changes are calculated using a pseudocount to avoid division by zero (**Methods**).

**Table S1-1.**
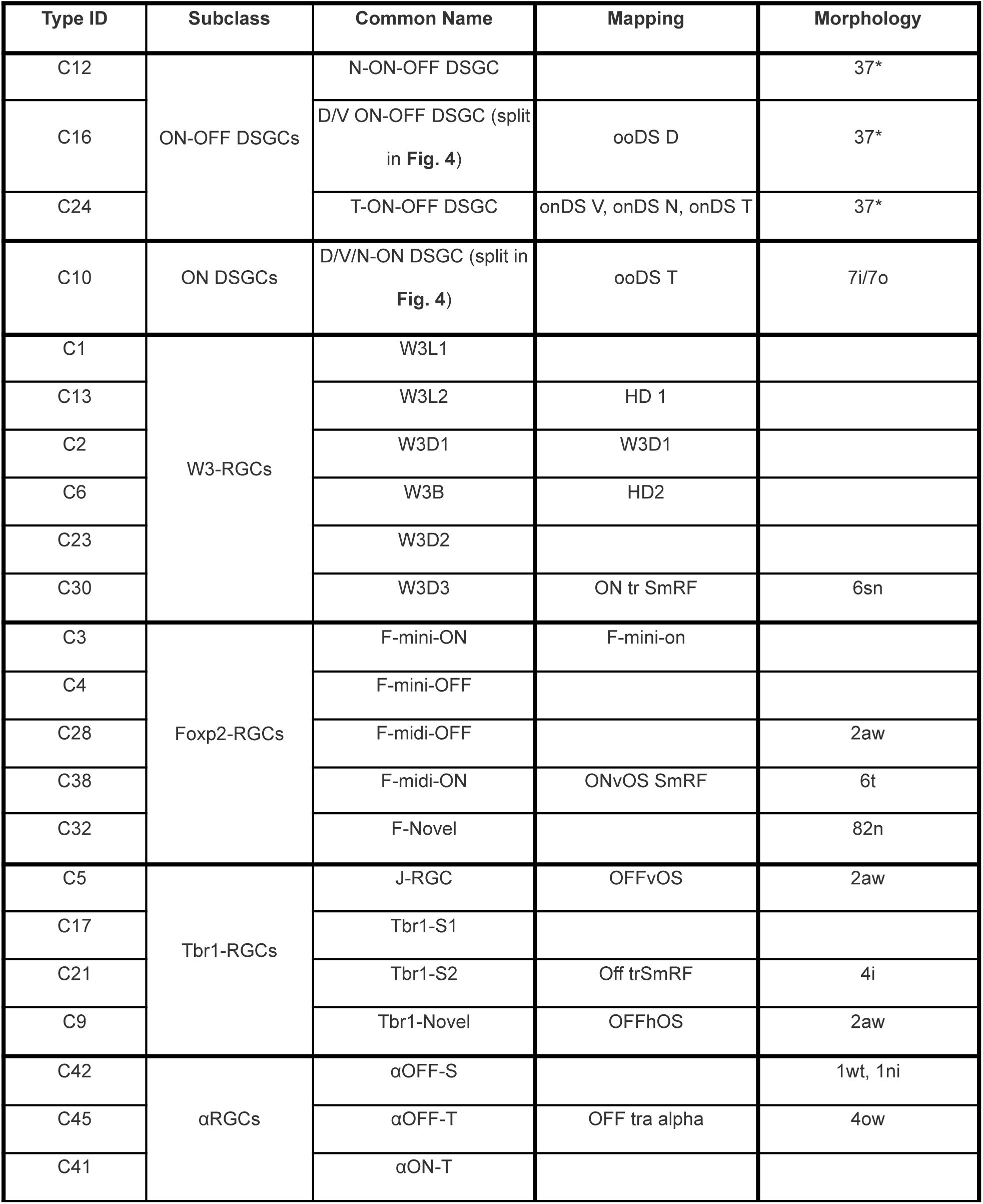
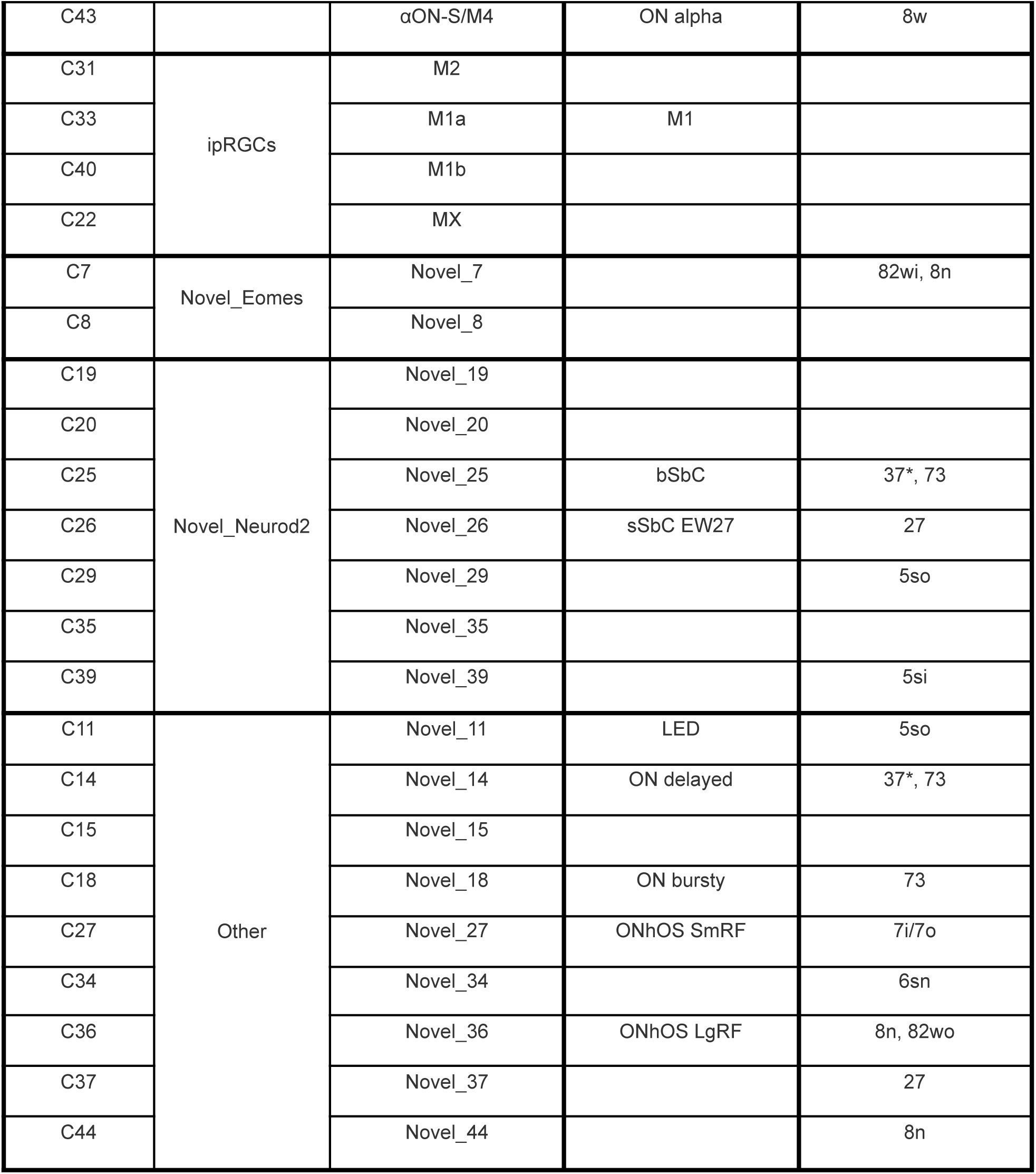
Common names associated with type IDs. Adapted from (Tran et al., 2019) with mappings from (Goetz et al., 2022) and morphologies from (Huang et al., 2022).

## Notes

Conflicts of interest: **None**

### Competing Interest Statement

The authors have declared no competing interest.

### Summary of Updates

New data added to support existing results

